# Molecular sterics guide tmRNA translocation on the ribosome

**DOI:** 10.64898/2026.10.08.757657

**Authors:** George Wanes, Kenneth C. Keiler, Christine M. Dunham, Paul C. Whitford

## Abstract

Translation of defective mRNA transcripts can lead to the formation of stalled “non-stop” ribosome complexes. To rescue these ribosomes, bacteria have evolved a mechanism called *trans*-translation that is mediated by transfer-messenger RNA (tmRNA) and small protein B (SmpB). During the translocation step of *trans*-translation, the stalled mRNA is ejected and the reading frame found in the mRNA-like domain (MLD) of tmRNA is loaded into the mRNA channel. The ribosome then translates the new template, thereby tagging the nascent protein for degradation. While structures of tmRNA-SmpB on the ribosome have been available for years, little is known about the molecular factors that govern the dynamics of tmRNA translocation. Here, we use an energetically smooth model (i.e. SMOG) to simulate translocation of tmRNA-SmpB, which reveals that the structure of the ribosome gives rise to a preferred sequence of conformational substeps. Specifically, the tRNA-like domain (TLD) translocates, then the MLD loads, and finally the C-terminal tail of SmpB rearranges. These steps are interdependent and associated with a range of sterically-induced barriers that regulate the kinetics. This provides a structural and energetic framework for understanding, and ultimately controlling, the dynamics of *trans*-translation.

## Introduction

During elongation a ribosome translates mRNA into protein until it encounters a stop codon. This process leads to release of the nascent protein and disassembly of the ribosome, allowing it to be recycled.^1^ However, this termination process can fail for a number of reasons, such as damage to transcripts, premature transcription termination or endonucleolytic mRNA cleavage. Erroneous frameshifting can also result in failure to recognize a stop codon. For these cases, the ribosome can translate the mRNA sequence until it reaches the 3’ end, resulting in a “non-stop” complex that is trapped and unable to terminate. ^2^ Since these ribosomes are non-productive, multiple rescue mechanisms have evolved in bacteria to ensure viability of the cell.^3,4^ The protein ArfA can recognize a non-stop complex and recruit release factor RF2,^5^ which allows the ribosome to terminate translation and disassemble. Similarly, ArfB can bind a non-stop ribosome, directly hydrolyze peptidyl-tRNA^6^ and release the nascent peptide. However, neither factor triggers degradation of the mRNA or incomplete nascent peptide. A more robust rescue mechanism is provided by tmRNA-SmpB: a ribonucleoprotein that simultaneously mimics the structure of tRNA and mRNA. It can bind the non-stop ribosome and introduce a tag to the nascent peptide that targets it for degradation by proteases,^7–10^ while the non-stop mRNA is degraded by RNase R.^11^ This is known as *trans*-translation, and it is the primary rescue mechanism in nearly all bacteria.^12^ Since there is not an analogous process in the human ribosome, it is a candidate target for broad-spectrum antibiotic development.^4^ Accordingly, understanding the mechanistic properties of *trans*-translation may allow for more effective therapeutic strategies to be developed.

tmRNA is a ~360-nucleotide RNA molecule and SmpB is a modest-sized (~ 150 residues) single-domain protein (Fig. 1). Each tmRNA molecule has a tRNA-like domain (TLD) that mimics the acceptor arm and elbow of tRNA (Fig. S1), and it is aminoacylated by alanyl-tRNA synthetase.^14^ The tag reading frame, also known as the mRNA-like domain (MLD), carries a short internal open reading frame that encodes the proteolysis tag. ^8^ In most bacteria, the two domains are connected by four pseudoknots and the H5 stem-loop, which together form a closed RNA ring.^15^ In some cyanobacteria and Alphaproteobacteria, tm-RNA is comprised of two separate RNA molecules: one contains the 5’ end and reading frame, while the other contains the 3’ end. ^16^ Upon reaching the so-called “accommodated” state, the TLD binds the A site of the large subunit (LSU), where SmpB substitutes for the missing mRNA on the small subunit (SSU; Fig. 1A-B).^13,17,18^ After tmRNA accommodation, the translocation step allows the ribosome to exchange templates. That is, the non-stop mRNA is released and translation resumes on the MLD sequence. During tmRNA translocation, the TLD moves between the A site and the P site of the ribosome, the MLD loads in the mRNA channel, and the resume codon is positioned within the decoding center^19^ (Fig. 1C-D). In addition, there are other large-scale conformational changes, including repositioning of the pseudoknot ring, displacement of H5 and rotation of the SmpB tail^13,18,19^ (Fig. S2). High-resolution cryo-EM structures have captured the endpoints of this transition (i.e. accommodated and post-translocation) states. ^13^ However, the precise sequence of substeps and physical factors/interactions that determine the dynamics have yet to be identified.

**Figure 1.**
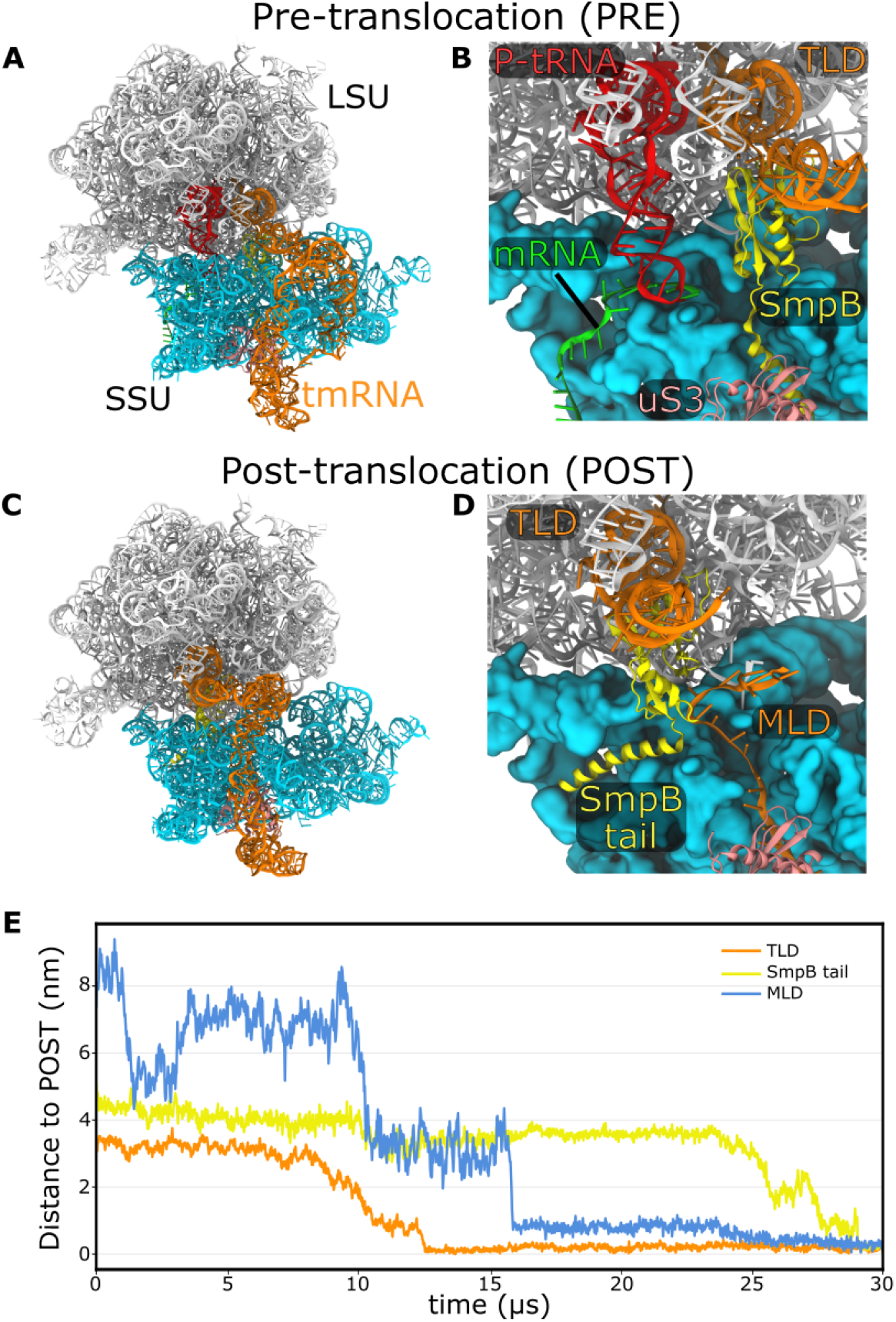
Translocation of tmRNA and SmpB between the ribosomal A and P sites. In the pre-translocation (PRE) state (panels A and B), the TLD is accommodated in the A site, while SmpB engages the decoding center, with its C-terminal tail occupying the mRNA entry channel (PDB ID: 7AC7^13^). The 23S rRNA, 16S rRNA, P-site tRNA, tmRNA, SmpB, uS3, and the mRNA are shown in white, cyan, red, orange, yellow, pink, and green, respectively. In the post-translocation (POST) conformation (panels C and D), the TLD and SmpB body have moved into the P site, with the SmpB tail rotated into the mRNA exit path, while the MLD loads into the mRNA channel and the A-site finger contacts pseudoknot 1 (PK1) of the tmRNA ring (PDB ID: 7ACJ^13^). (E) Representative simulated trajectory of tmRNA translocation using an all-atom structure-based (SMOG) model. Time traces of different components reveal a distinct order of events. First, the TLD advances toward the P site almost monotonically. Next, the MLD shows reversible fluctuations and pauses at multiple intermediate positions before settling into the mRNA channel, pointing to several steric obstacles along its path. Finally, the SmpB tail adopts its POST position, which requires repeated attempts before reaching the E site, reflecting the presence of a pronounced barrier.

A strategy that has been effective for probing how molecular structure guides dynamics is the application of molecular dynamics simulations with all-atom structure-based “SMOG” models.^20^ In this class of models, experimental structures are explicitly defined as minima on the energy landscape. By providing a smooth description of the global energetics, these models are well suited to identify sterically-induced barriers that arise from imperfect complementarity of ribosomal components.^21^ Further, since molecular flexibility is well described by these models,^22^ they can indicate whether configurational entropy will favor specific conformational pathways. There are many examples where these models have implicated sterically- and entropically-induced barriers during normal translation. For tRNA-mRNA translocation, they predicted the presence of SSU head tilting at a late stage of translocation.^21^ By introducing perturbations to the model, it was demonstrated how tilting arises from specific steric interactions between the tRNA and ribosome. In support of this sterics-based approach, the predicted tilting motion was consistent with subsequent single-molecule measurements.^23,24^ When applied to tRNA P/E hybrid state formation, SMOG models similarly showed how a constriction can limit the kinetics. ^25^ These models have been applied to many other molecular assemblies, such as viral fusion proteins,^26^ where a balance of disorder and steric factors dictates the structural dynamics. In all cases, the simplified character of the models has allowed for identification of structural features that control the kinetic and mechanistic properties of large-scale conformational rearrangements.

In the current study, we use SMOG models to determine whether molecular structure/sterics predispose tmRNA-SmpB to access specific conformational pathways during translocation, and to identify the molecular features that most strongly influence the dynamics. By simulating translocation using multiple variants of the model, we show that the molecular architecture of the ribosome and tmRNA lead to a well-defined sequence of conformational substeps. Despite the simplified character of the models, tmRNA-SmpB translocation is associated with a complex landscape and a range of intermediate states. These sterically-induced and entropically-favored intermediates arise directly from the structure, suggesting that tmRNA-SmpB has evolved to enforce a specific order of conformational events, lending robustness to the rescue process.

## Results

### Simulating tmRNA translocation on the ribosome

Before describing how SMOG models were applied to tmRNA translocation, it is worthwhile to briefly discuss some key features of the models. First, all non-H atoms are explicitly represented, which ensures that the steric composition of the complex is accurately described. Next, specific experimental structures are predefined to be minima on the potential energy landscape. The rationale for this assignment is that stable experimentally-resolved structures are necessarily free-energy minima. Since a complete steric description is applied, these models are suited to determine whether the sterics/structure impose strong limits on the conformational pathways, and to identify the relative contributions of each structural element. The current study uses a SMOG model in which the ribosome and tmRNA have competing energetics. For the ribosome, the classical conformation is explicitly defined to be stable. This assignment is consistent with the observation that a non-stop ribosome will adopt a classical conformation with a tRNA molecule in the P/P conformation (PDB IDs: 7ABZ, 7AC7^13^). All contacts with tmRNA-SmpB are defined to stabilize the post-translocation (POST) structure. With this model, tmRNA translocation is described as a process where tmRNA induces the conformational changes in the ribosome that are required for template switching. For complete model details, see methods.

Simulations of tmRNA translocation with the SMOG model reveal a complex process involving sequential conformational motions of tmRNA and SmpB (Supplementary Movie 1). To identify the order of events, we defined three reaction coordinates: *R*_TLD_, *R*_MLD_, and 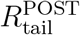 (Fig. S3). Each is a distance from the POST conformation, calculated after aligning the LSU rRNA. *R*_TLD_ describes movement of the TLD (U9). *R*_MLD_ monitors the first nucleotide of the resume codon (G90). 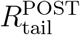 is the distance of the C-terminal tail of SmpB (Glu146). See Methods for complete list of coordinates used in this study.

In a representative time trace (Fig. 1E), the TLD reaches the POST configuration first, and it appears to gradually progress (along with the body of SmpB) towards the ribosomal P site in a nearly monotonic fashion. As expected, due to the strong binding of SmpB to tmRNA via its oligonucleotide-binding fold,^27^ movement of the globular domain of SmpB (i.e. the “body”) is highly correlated with TLD movement (Fig. S4). In contrast, the MLD is found to undergo large-scale reversible fluctuations towards the mRNA channel. In this specific simulation, the MLD (resume codon) approaches the mRNA channel and reaches a distance of ~ 5 nm, before returning to more distant positions. Later, as the TLD approaches the P site, the MLD again fluctuates towards the mRNA channel and samples distances of ~ 3 *−* 4 nm for an extended period of time, consistent with the presence of a sterically-induced intermediate. After the TLD loads, the MLD proceeds further and fully associates with the mRNA channel (*R*_MLD_ ~ 0.2 nm). In order to complete the transition to the POST state, the SmpB tail makes multiple attempts and eventually associates with the E site of the SSU. As described above, due to the simplicity of this model, the presence of intermediates and a specific ordering of events suggests that the steric composition of the ribosome determines the mechanism and kinetics of tmRNA translocation.

### Structure dictates the mechanism of tmRNA translocation

To characterize the range of conformational pathways that may be accessed during translocation, we performed 100 independent simulations in which tmRNA-SmpB transitioned between PRE (Fig. 1A,B) and POST (Fig. 1C,D) conformations. Consistent with the representative trajectory (Fig. 1E), this set of simulated transitions indicates that steric interactions strongly favor pathways in which the TLD translocates prior to the MLD. Specifically, in 98 (of 100) simulations, the TLD was found to associate with the P site of the LSU prior to MLD association with the mRNA channel. To describe the full set of simulated events, we calculated the probability as a function of *R*_TLD_ and *R*_MLD_ (Fig. 2A). There are multiple notable features in this distribution. First, as expected, the dominant pathway involves sequential movement of the TLD and MLD, where the TLD reaches the P site (*R*_TLD_; S 0.2 nm) while *R*_MLD_ is still large (~6 nm; Fig. 2C). This suggests that TLD translocation may serve to prepare the MLD for loading. This TLD-dependence of the MLD may be rationalized in terms of the structure. Since SmpB remains bound to the TLD throughout translocation, movement of the TLD necessarily leads to displacement of the SmpB tail (Fig. S5), which is required for the MLD to associate.

**Figure 2.**
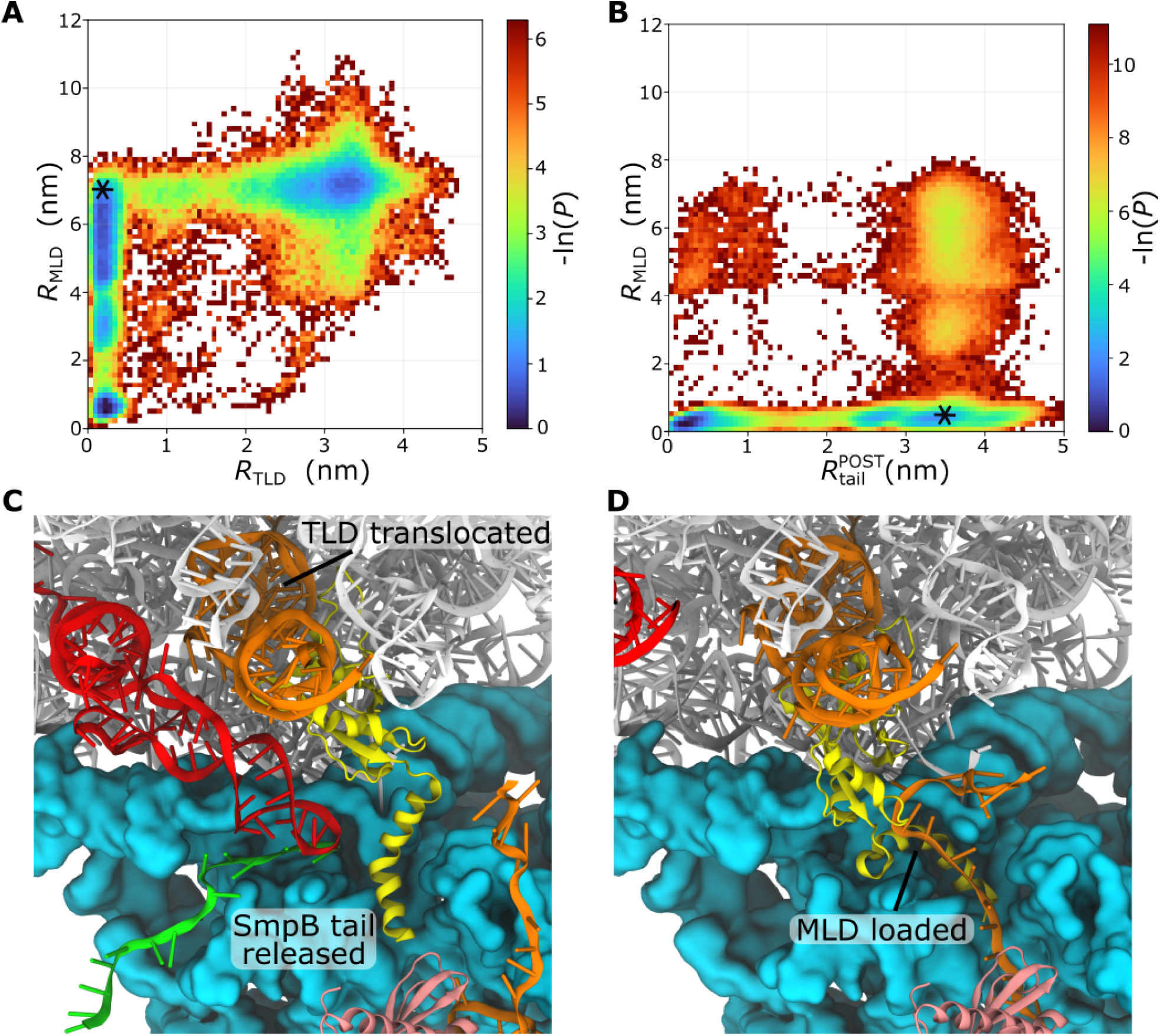
Structural properties of tmRNA translocation. (A) The 2D probability distribution as a function of the distances of the TLD (*R*_TLD_) and MLD (*R*_MLD_) from their POST positions reveals that, along the dominant pathway (starred), the TLD translocates prior to MLD loading into the mRNA channel. (B) 2D conditional probability distribution of *R*_MLD_ and 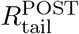 (distance of SmpB tail from POST position), calculated for *R*_TLD_*<* 0.5 nm (i.e. TLD translocated). The distribution shows that MLD loading into the mRNA channel typically precedes full translocation of the SmpB tail (starred), where the SmpB tail is partially withdrawn from the mRNA channel. (C) Representative structure with the TLD in the P site and the SmpB tail partially separated from the mRNA channel. (D) Representative structure with TLD and MLD in their POST positions while the SmpB tail is near the A site.

Even though the model defines the POST conformation as stable, PRE-like conformations (*R*_TLD_ ~ 3 nm; Fig. 2A) also appear to be stabilized by multiple steric features. First, the N-terminal tail of bL27, which is adjacent to the peptidyl transferase center, can impede TLD movement (Fig. S6) and lead to prolonged sampling of the PRE/accommodated state.^28^ Next, the P-site tRNA sterically occludes the P site until it is released from the ribosome (Fig. S7). Finally, steric interactions with the A-site finger (ASF; helix 38 of the 23S rRNA) impede TLD displacements, further amplifying the sampling of the PRE/accommodated ensemble (Fig. S8).

In addition to predicting that TLD movement occurs early in the translocation process, the model also indicates that the structure of the ribosome favors MLD loading prior to binding of the SmpB tail to the E site (Fig. 2B,D). To quantify this, we calculated the probability as a function of 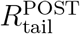 and *R*_MLD_ (Fig. 2B). MLD and SmpB tail movement are both associated with multiple intermediate ensembles. However, there is variability in the relative timing of MLD and SmpB tail rearrangements. MLD loading usually precedes SmpB tail docking, though the tail occasionally only reaches intermediate/near-POST positions prior to completion of MLD loading. As described later, in some simulations the tail appears to become trapped and does not fully adopt the POST position.

To confirm that the observed mechanistic features arise from structure, rather than the details of the energetics, we compared the dynamics when using variants of the SMOG model (Tab. S1). In the first variant, the strength of the tmRNA-SmpB-ribosome interactions was increased, or decreased, to test whether changes in timescales impacts the mechanism. SSU head flexibility was also decreased to assess whether more rapid MLD loading could lead to a change in the relative timing of TLD and MLD rearrangements. ASF flexibility was perturbed to determine whether changes in TLD kinetics impacts the overall mechanistic properties. Finally, the affinity of tRNA and mRNA for the ribosome was varied to determine whether P-site stability influences the mechanism. While each change represent a significant change to the energetics (up to tens of *k*_B_*T*), we find consistent ordering of the TLD, MLD and SmpB tail rearrangements with all models (Fig. S9,S10). This robust character highlights how the structures of the ribosome and tmRNA-SmpB impose stringent limitations on the accessible pathways. Together, they enforce a specific sequence of conformational transitions that begins with displacement of the TLD.

### MLD loading involves multiple barriers

We next asked whether steric interactions between tmRNA and the ribosome impact loading of the MLD. For this, we describe the MLD in terms of two regions: the resume codon (G90) and uS3-binding segment (A97-A103; Fig. S2). Given their relative positions in sequence, we refer to the uS3-binding segment as the downstream MLD (dMLD), while the term MLD will describe the resume codon. The SMOG model implicates a strong preference for the dMLD to associated with uS3 (Fig. S11A) before the resume codon reaches the A site. As shown below, there is an interplay between these two steps, where dMLD binding leads to more rapid MLD loading.

After the dMLD binds uS3, the MLD transits across a sequence of sterically-induced barriers. To identify these barriers, we calculated the probability as a function of *R*_MLD_ (shown as the negative logarithm in Fig. 3A). With this coordinate, barriers are apparent at *R*_MLD_ ~3.8 nm and ~1.5 nm (Fig. 3A). As the MLD crosses each barrier, it forms distinct steric interactions with the ribosome. For the first barrier (*R*_MLD_ ~ 3.8 nm), the MLD frequently contacts h18 (U531-A532, C513-C514) in the SSU body, and h34 (m^2^G1207-C1208) in the head (Fig. 3B and Fig. S12A). Similarly, at the second barrier the MLD contacts G530, A1196, C1054 and C1397, also found in the SSU body and head (Fig. 3C and Fig. S12B). On a technical note, it is important to qualify our identification of these barriers. Since the simulated trajectories do not reflect equilibrium sampling (i.e. terminated upon reaching POST), the distributions cannot directly describe the free energy. However, this provides a steady-state distribution,^22^ which can reliably identify signatures of freeenergy barriers.^29^ With this in mind, the simulations provide evidence that MLD loading is associated with sterically-induced free-energy barriers.

**Figure 3.**
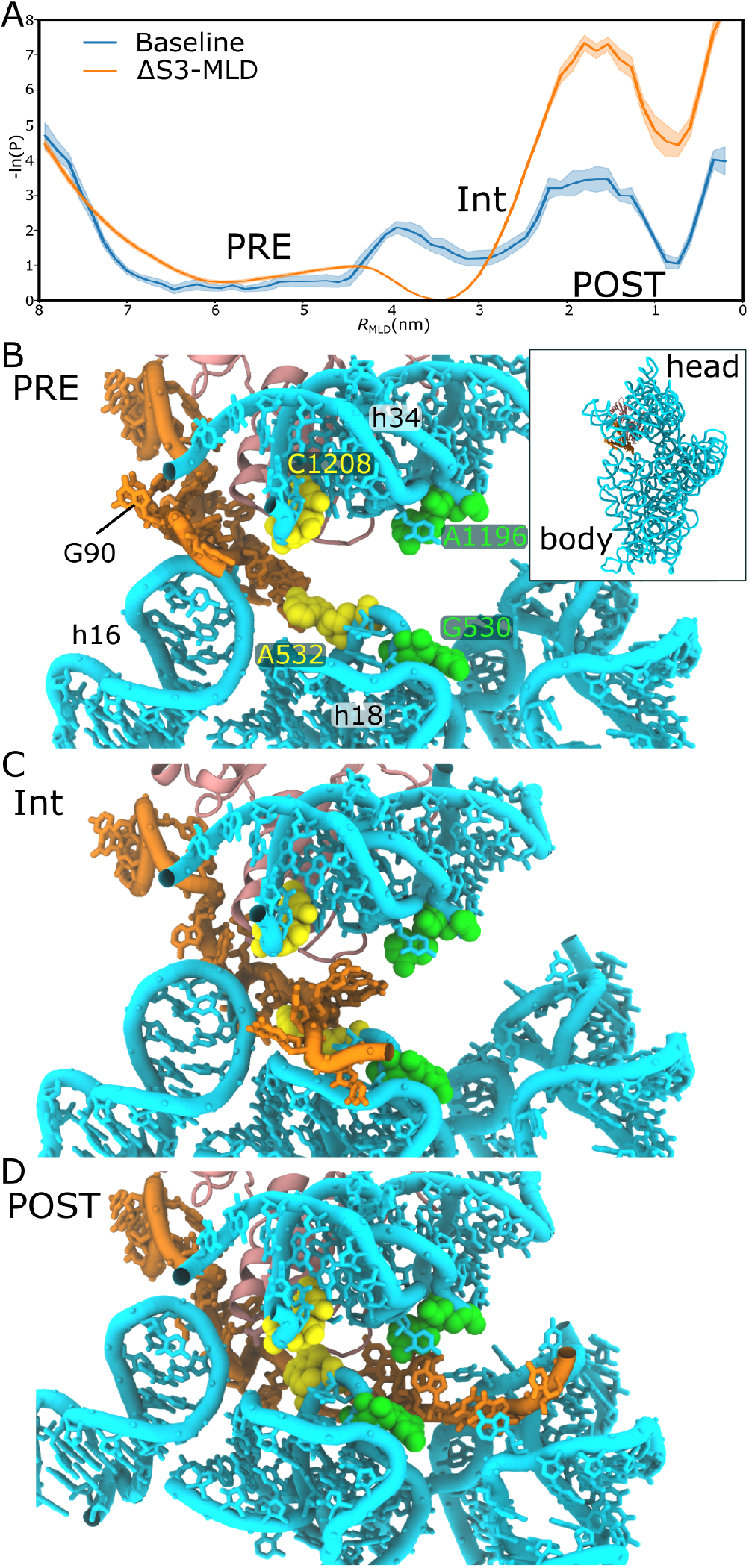
MLD loading proceeds through multiple steps. (A) The probability distribution as a function of *R*_MLD_ computed from frames satisfying *R*_TLD_ *<* 0.5 nm (i.e. TLD translocated), including all frames until *R*_MLD_ adopts a value below 0.5 nm. The distribution shows that the MLD progresses across multiple steric barriers, dwelling at ~6 nm and ~3 nm before fully loading into the mRNA channel. As shown in Fig. S11, downstream residues of the MLD form contacts with protein uS3 at the mRNA entry channel, which help the MLD load. When the downstream contacts are removed (ΔS3-MLD), there is a significantly higher barrier preceding the fully loaded state. Shaded regions represent the standard error of the mean at each point along the coordinate, estimated by grouping the 100 independent trajectories into 10 blocks of 10 trajectories each and computing the distribution independently for each block. (B) Structural representations of the MLD at ~6 nm from its loaded position, sterically impeded by one extension of the 30S latch, via residues h18 (U531-A532, C513-C514) and h34 (m^2^G1207-C1208; Fig. S12A). (C) The MLD is obstructed at ~3 nm by another extension of the latch, contacting G530 from the body and A1196, C1054 and C1397 from the head (Fig. S12B). Once it crosses this barrier, it can reach the mRNA channel, as shown in (D).

The sequential association of the dMLD and MLD suggests they are coupled during loading. To explore this relationship, we constructed another variant of the SMOG model. In this SMOG model variant (ΔS3-MLD model; Tab. S1), all stabilizing interactions between the dMLD and uS3 are removed. We find that removal of these interactions amplifies the barrier separating the intermediate and POST states (Fig. 3A). This is consistent with a roughly 12-fold increase in the loading time (time required to reach *R*_MLD_ *<* 0.5 nm; Fig. S11C). These signatures strongly suggest that MLD loading is a multi-step process, where early formation of uS3 interactions facilitates proper positioning of the resume codon (Fig. S11A,B). These calculations also implicate a dynamic interplay between the dMLD, pseudoknot 1 (PK1) and MLD. Upon translocation of the TLD, PK1 forms stabilizing contacts with the ASF (Fig. S13). In the simulations, PK1-ASF interactions localize PK1, allowing it to serve as an anchor point that tethers the MLD to the ribosome. Since PK1 is upstream of the MLD (Fig. S2) and the dMLD is downstream, dMLD docking can naturally “tighten” the single-stranded MLD. This tightening localizes the MLD, which can help it navigate the sterically-dense environment imposed by the SSU.

There are multiple prior observations that suggest dMLD-uS3 interactions are rather strong, which would be a required for the predicted dMLD-MLD relationship to manifest. In the POST structure (PDB ID: 7ACJ), the dMLD (specifically G99 and A101) forms close interactions with positively charged residues on uS3 (Arg127, Arg132, and Arg136), where Arg132 interacts with both G99 and A101 of the tmRNA (Fig. S11D). There is also evidence that uS3 has helicase activity that unwinds the secondary structure of mRNA in bacteria^30,31^ or stabilizes mRNA contacts with the pre-initiation complex in eukaryotes. ^32^ For either of these roles, uS3 would need to be capable of strongly binding single-stranded RNA. Here, we show how these relatively strong interactions could also influence the kinetic and mechanistic properties of tmRNA translocation.

### SmpB mobility can stall translocation

tmRNA translocation requires the C-terminal helix of SmpB to undergo a ~ 50 Å displacement in order to bind the E site of the mRNA channel on the SSU (Fig. 1B,D). We find that this rearrangement may occur via two distinct paths, which we will refer to as “direct” and “indirect”. To quantify the dynamics of SmpB, we use the coordinates 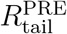 and 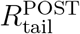: the distances of the C-terminal helix (C_*α*_ atom of Glu146) from the PRE and POST positions. In the direct pathway, 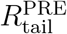 and 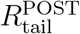 are anticorrelated, which would be associated with paths that minimize the distance traversed by the tail (Fig. 4B,C). In contrast, indirect transitions are associated with simultaneous increases in both distances at intermediate time points (Fig. 4D,E).

**Figure 4.**
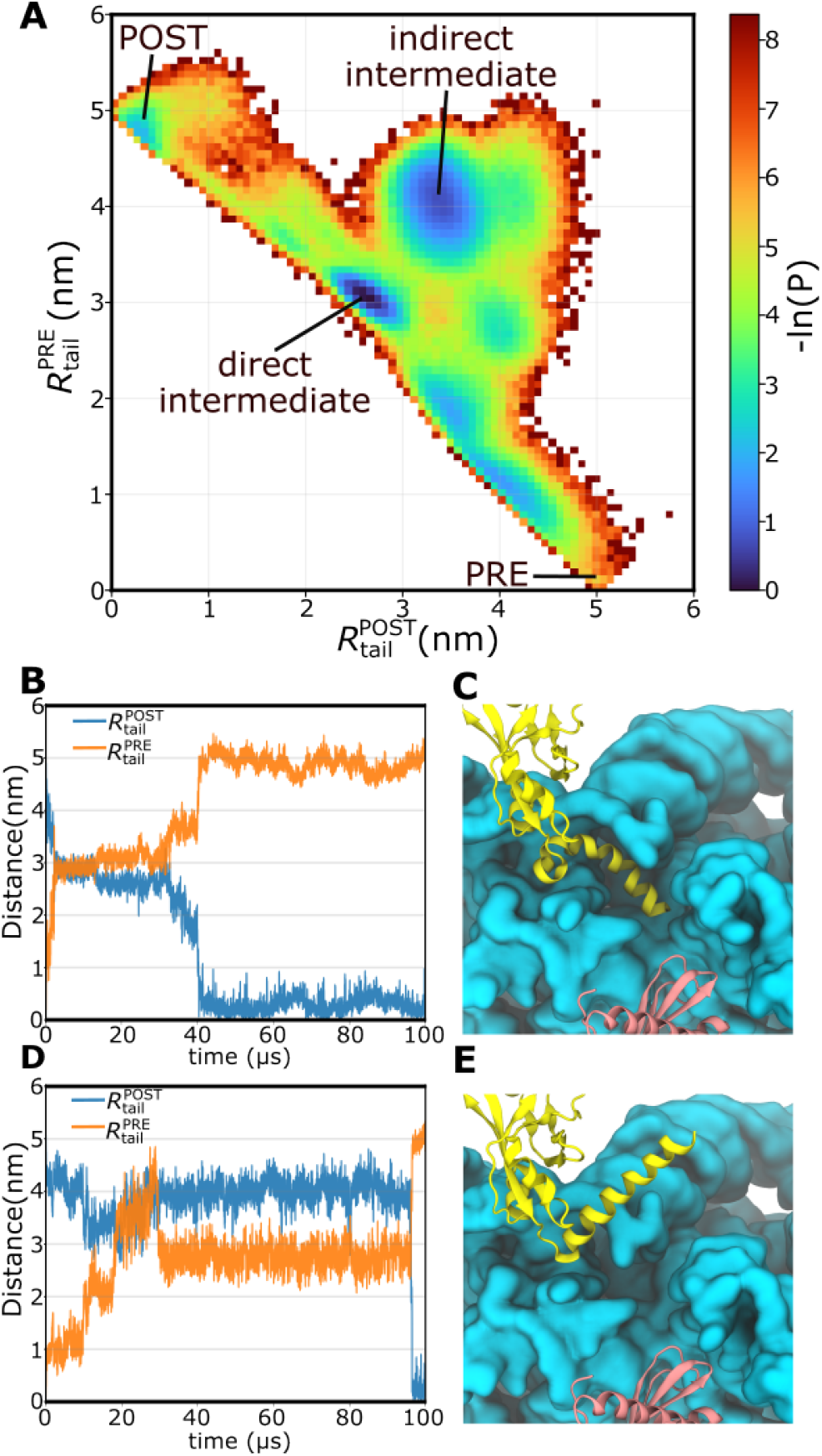
The SmpB tail encounters steric and entropic barriers during translocation. (A) The probability density as a function of the SmpB tail distance from the initial 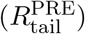 and final 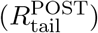 positions reveals multiple intermediate states: a direct intermediate, along the dominant translocation pathway, and an indirect intermediate which represents an off-path intermediate. (B) Representative time trace of 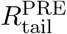 and 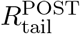 for the direct route where the coordinates are anticorrelated, signifying displacement away from the PRE position and toward the POST position. (C) Representative structure of SmpB in the direct intermediate. The SmpB body has translocated while the tail remains partially confined near the mRNA channel. (D) Representative time trace of 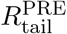 and 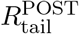 for the indirect pathway, where the two coordinates simultaneously increase at intermediate time points. (E) The indirect intermediate represents an off-path state in which the tail rotates away from the mRNA channel and its POST position.

We find evidence of multiple intermediates along the direct and indirect paths (Fig. 4A). The dominant intermediate associated with the direct path is characterized by interactions with residue m^4^Cm1402, C1403 and m^3^U1498 of the SSU (Fig. S14). Specifically, contacts are formed in more than 90% of the simulated conformations in the direct intermediate (2.5 nm *<* 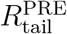 *<* 3.5 nm and 2 nm *<* 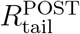 *<* 3 nm), revealing the presence of a significant steric hindrance. In contrast to the direct intermediate, the indirect intermediate is associated with a more dispersed set of configurations. While the C-terminal end of the tail (Trp147-Arg160) is generally positioned in the vicinity of helix 44, it does not consistently contact specific residues in the ribosome. Instead, the end of the SmpB tail appears to be solvent exposed, consistent with a larger contribution of configurational entropy in this state.

The presence of direct and indirect intermediates suggests a potential role of EF-G during tmRNA translocation. In our simulations, the observed dwell time in the indirect intermediate is at least twice as long as that of the direct intermediate. Of the 40 simulations that sampled the indirect intermediate, there were 13 in which the tail did not escape. Since the tail did not reach POST in some of the indirect events, the ratio of indirect and direct timescales is a lower bound. Interestingly, the position of the tail in the indirect ensemble overlaps with the location of domain IV of EF-G in the post-translocation state of tmRNA-SmpB^19^ (Fig. S15). If EF-G engages the ribosome prior to the completion of tmRNA translocation, one would expect that the indirect intermediate would be sterically prohibited by domain IV. This would be consistent with biochemical analysis that found EF-G accelerates the rate of mRNA dissociation during tmRNA translocation.^33^ That is, EF-G may prevent the SmpB tail from becoming kinetically trapped, thereby facilitating access to the direct route and more rapidly displacing the non-stop mRNA.

## Discussion

Despite advances in structure determination, a complete description of tmRNA dynamics during ribosome rescue has remained elusive. Since tmRNA-based rescue is specific to bacteria, insights into the mechanistic properties may provide new opportunities for novel antibiotic development. To this end, we have focused on identifying and characterizing sterically-induced barriers that are associated with a large-scale rearrangement of tmRNA-SmpB on the ribosome. In future studies, it may be possible to target the predicted intermediates by binding small molecules that effectively amplify the steric interactions and arrest the dynamics.

In addition to potential therapeutic applications, the mechanistic framework established here provides a foundation that can guide new experimental strategies, while also rationalizing previous empirical observations. As an example, one may consider the predicted impact of PK1, where association with the ASF can help localize and load the MLD. Consistent with this, Valle et al.^34^ suggested that tmRNA interactions between PK1 and ribosomal elements might be needed to correctly position the resume codon for entry into the decoding site. Similarly, Nameki et al. ^35^ showed that mutations in PK1, particularly in the U60-G62 region, affect alanine incorporation by introducing a defect in translocation or trans-peptidation.

Finally, Tanner et al. ^36^ used *in vivo* genetic selection to score survival above a threshold as “active” without distinguishing degrees of activity. Their study found that PK1 could be replaced by unrelated hairpin sequences, though some retained only a small fraction of wild-type activity. Based on this, they concluded that stable secondary structure in this position, rather than the specific pseudoknot fold, is necessary for tmRNA to function. These disparate observations may all be understood in terms of the predicted PK1-mediated impact on MLD loading, where the only requirement is a nominal affinity of PK1 for the ASF.

In the coming years, it will be interesting to build upon the current study and partition the full range of molecular factors that contribute to tmRNA dynamics on the ribosome. While the current study identifies a range of sterically-induced barriers, these interactions often involve RNA-RNA interactions. Accordingly, the precise free-energy barriers associated with tmRNA movement will be sensitive to ionic conditions. In future studies, SMOG models that include ionic interactions^37,38^ will likely prove useful to establish the precise balance between steric and electrostatic interactions. In addition to electrostatics, there are still many questions regarding the exact role of EF-G during tmRNA translocation. Beyond translocation, there are other large-scale conformational changes, such as accommodation, where each is likely to be associated with a complex interplay of molecular components. Through the continued development of novel theoretical methods, combined with advanced experimental techniques, the field is positioned to establish a comprehensive understanding of *trans*-translation.

## Methods

### Structure-based “SMOG” model

To investigate translocation of tmRNA-SmpB on the ribosome, we use structure-based models where all non-hydrogen atoms are explicitly represented. Biomolecular interactions are defined to stabilize preassigned structures.^20^ Since the free-energy minima arise from the full energetic composition of the system, which includes electrostatic and solvation effects, the SMOG model describes an effective potential energy. ^39^ That is, the potential energy implicitly describes the contribution of all factors that give rise to the experimental structure. While the current study focuses primarily on sterically-induced barriers, it is important to note that these models can be extended to explicitly account for additional energetic factors, such as ionic effects. ^38^ The functional form of the potential energy used in this study may be expressed as:

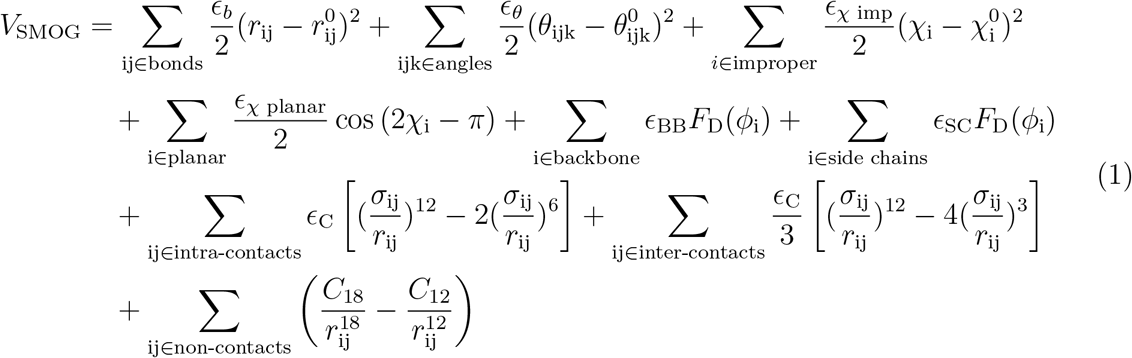

where

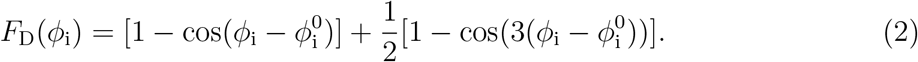

*V* _SMOG_ describes the inter- and intramolecular interactions of the biomolecule. Here, intra-ribosomal contacts are defined to stabilize the PRE/accommodated conformation (PDB 7AC7^13^) and tmRNA-SmpB interactions stabilize the POST state (PDB 7ACJ^13^). Bonded parameters 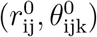 are assigned values given in the Amber99sb-ildn force field. ^40^ To avoid artificial expansion of the system, ^21^ contact distances (*σ*_ij_) are set to 0.96 of the value found in the experimental structure. Energetic weights are set as following 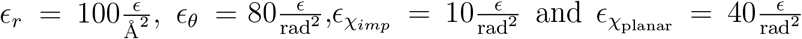 with *ε* denoting the reduced energy unit. To define the contacts, Shadow Contact Map algorithm^41^ is used with the default 6 Å cutoff and 1Å shadowing radius. Intramolecular contacts within the tmRNA–SmpB complex and the ribosome are described using an attractive 6–12 potential. Additionally, the contacts between the tmRNA–SmpB and the ribosome were modeled with a long-range 3–12 potential with the stable minimum assigned from the POST state. The contacts and dihedrals weights are normalized as previously described.^20^ The 12-18 term represents the excluded volume for atom pairs that are not in contact in the experimental structure. The C_18_ and C_12_ parameters are given values to mimic the repulsive component of 6-12 interactions in Amber99sb-ildn force field,^40^ as described previously.^37^

For structure-based model, stabilizing contacts describe effective interactions. ^39^ Contacts between ribosome and tmRNA-SmpB were rescaled by a factor of 0.6, to reflect the transient nature of the interactions, relative to contacts that maintain the structural integrity of the ribosome. Additionally, contacts between the 30S ribosomal subunit and the A-site finger (ASF) were rescaled by 0.2, allowing for mobility of the ASF, consistent with many crystallographic and cryo-EM structures. Contacts between the body and head of the small subunit and between the head of the small subunit and the large subunit were rescaled by a factor of 0.3 to allow for the head tilting and rotation. In describe post-peptidyltransfer state, a harmonic potential with weight 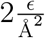 was introduced between the P site of the ribosomal large subunit (phosphate atom of C2064) and the 3’-CCA tail of the tmRNA (phosphate atom of A363). The overall mechanistic properties were robust to the precise strengths of these sets of contacts (Fig. S9).

### Simulation details

Initial force field files were generated using SMOG 2^42^ with the AA 12-18 Wang22.v1 templates.^37^ Further manual modifications were introduced to the force field files to include the 3-12 potential. Simulations were performed using OpenSMOG v1.2^43^ and OpenMM 8.^44^ Simulations were performed using Langevin dynamics at a reduced temperature of 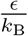 with a timestep of 0.002. This temperature was selected because the flexibility it yields is consistent with explicit-solvent simulations and crystallographic B-factors of molecular assemblies.^45^ In order to estimate the effective simulated time, we set one reduced time unit equal to roughly 1 ns. This estimate was previously derived for tRNA movement within the ribosome,^46^ based on comparison of the diffusion coefficients obtained from SMOG and explicit-solvent simulations.

### Coordinates used for analysis

To characterize the simulated dynamics, the following coordinates were used, each defined as the distance of a reference atom from its position in the POST state (unless otherwise noted), after alignment on the LSU rRNA:

- *R*_TLD_ - Describes movement of the TLD. Defined by the P atom of U9.
- *R*_MLD_ - Describes movement of the MLD (resume codon). Defined by the P atom of G90.
- *R*_body_ - Describes movement of the body of SmpB. Defined by the C_*α*_ atom of Leu65.
- 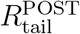 - Describes movement of the tail of SmpB. Defined by the C_*α*_ atom of Glu146.
- 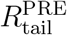 - Describes movement of the tail of SmpB, defined by the same atom as 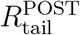 (C_*α*_ of Glu146), but relative to the PRE state.

## DATA AVAILABILITY

Upon publication, all input decks and analysis scripts will be released on github and registered with Zenodo. Due to the size of the data set, it is not feasible to deposit all simulated trajectories in a public repository. Accordingly, representative simulated trajectories will be made available on Dryad and the full set of trajectories will be available upon reasonable request.

## ACKNOWLEDGMENTS

This work was supported by the National Institutes of Health and the National Science Foundation.

## AUTHOR CONTRIBUTIONS

G.W. performed all simulations and conducted formal analysis. All authors contributed to the design of the study, interpretation of the data and manuscript preparation.

