## Supplementary Methods and Figures for "Molecular sterics guide tmRNA translocation on the ribosome"

#### Contents

**1 SI Methods** **S2**

#### List of Figures

S1 The tRNA-like domain (TLD) of tmRNA mimics the tRNA acceptor arm . . . . . . . . . . . S4
S2 Structure of tmRNA on the ribosome . . . . . . . . . . . . . . . . . . . . . . . . . . . . . . S5
S3 Coordinate definitions . . . . . . . . . . . . . . . . . . . . . . . . . . . . . . . . . . . . . S6
S4 SmpB body and TLD movements are coupled . . . . . . . . . . . . . . . . . . . . . . . . . S7
S5 TLD translocation leads to release of the SmpB tail from the mRNA channel . . . . . . . . . S8
S6 The N-terminal tail of bL27 impedes TLD translocation . . . . . . . . . . . . . . . . . . . . S9
S7 The P-site tRNA sterically impedes TLD translocation . . . . . . . . . . . . . . . . . . . . . S10
S8 The A-site finger (ASF) can hinder TLD motion . . . . . . . . . . . . . . . . . . . . . . . . S11
S9 Robustness of the translocation mechanism across model variants. . . . . . . . . . . . . . . . S12
S10 Robustness of the translocation to the affinity of tRNA/mRNA for the ribosomal P site . . . S13
S11 dMLD–uS3 interactions facilitate MLD loading . . . . . . . . . . . . . . . . . . . . . . . . S14
S12 SSU residues that contact MLD during loading . . . . . . . . . . . . . . . . . . . . . . . . S15
S13 PK1 forms contacts with the ASF upon TLD translocation. . . . . . . . . . . . . . . . . . . S16
S14 SSU residues interact with the SmpB tail in the direct intermediate . . . . . . . . . . . . . . S17
S15 EF-G overlaps with the position of the SmpB tail in the indirect intermediate . . . . . . . . . S18

### 1 SI Methods

In addition to the primary model that was used (i.e. the baseline model), we also constructed a number of variants of the model in order to assess the robustness of the mechanistic features and to establish direct relationships between different molecular components. The following models were constructed:

- Baseline - This is the primary model that was used to simulate tmRNA-SmpB translocation. Main methods describe the full details of this model.
- Weaker binding - In this model, the tmRNA and SmpB contacts with the ribosome were reduced in strength, to assess the robustness of the translocation mechanism. Results are shown in Figure S9.
- Stronger binding - In this model, the tmRNA and SmpB contacts with the ribosome were increased in strength, to assess the robustness of the translocation mechanism. Results are shown in Figure S9.
- Less flexible SSU head - In this model, the contacts between the SSU head and the LSU/SSU body were stronger, relative to the baseline model. This model was used to assess whether MLD loading kinetics could lead to a change in the relative timing of TLD and MLD rearrangements. Results are shown in Figure S9.
- Less flexible ASF - In this model, the contacts between the ASF and the SSU were stronger, relative to the baseline model. This model was used to determine whether TLD translocation kinetics affects the overall mechanistic properties. Results are shown in Figure S9.
- More flexible ASF - In this model, the contacts between the ASF and the SSU were weaker, relative to the baseline model. This model was used to determine whether TLD translocation kinetics affects the overall mechanistic properties. Results are shown in Figure S9.
- tRNA affinity v1 - In this model, the contacts between the tRNA/mRNA with the ribosomal P site were stronger, relative to the baseline model. This model was used to determine whether tRNA interactions with the ribosomal P site affect the overall mechanistic properties. Results are shown in Figure S10.
- tRNA affinity v2 - In this model, the contacts between the tRNA/mRNA with the ribosomal P site were stronger, relative to the baseline model. This model was used to determine whether tRNA interactions with the ribosomal P site affect the overall mechanistic properties. Results are shown in Figure S10.
- ΔS3-MLD - In this model, the contacts between the downstream MLD (dMLD; A97-A103) and the uS3 protein were removed. This model was used to determine whether dMLD contacts with uS3 affect MLD loading mechanism and kinetics. Results are shown in Figures 3 and S11
- ΔASF-TLD : The contacts in this model are identical to the baseline model. However, the excluded volume term (12-18 term in Eq. 1) between the TLD and the ASF residues is switched off. This model was introduced to investigate the steric effects of the ASF on the TLD's progression to the P site. Results are shown in Figures S8.

The full set of model parameters and number of replicas simulated are given in Table S1.

**Table S1:** Contact strengths used for the Baseline model and all variants. Contact weights in the table correspond to the scaling factor of each group of contacts, relative to the default weight [1].

| Model | tmRNA-SmpB–Ribosome | Head–Body/LSU | SSU–ASF | tRNA/mRNA-P site | dMLD-uS3 | # of replicas |
|---|---|---|---|---|---|---|
| Baseline | 0.6 | 0.3 | 0.2 | 0.0 | 0.6 | 100 |
| Weaker binding | 0.4 | 0.3 | 0.2 | 0.0 | 0.6 | 40 |
| Stronger binding | 1.0 | 0.3 | 0.2 | 0.0 | 0.6 | 40 |
| Less flexible SSU head | 0.6 | 0.5 | 0.2 | 0.0 | 0.6 | 100 |
| Less flexible ASF | 0.6 | 0.3 | 0.4 | 0.0 | 0.6 | 40 |
| More flexible ASF | 0.6 | 0.3 | 0.0 | 0.0 | 0.6 | 40 |
| tRNA affinity v1 | 0.6 | 0.3 | 0.2 | 0.1 | 0.6 | 8 |
| tRNA affinity v2 | 0.6 | 0.3 | 0.2 | 0.2 | 0.6 | 8 |
| ΔS3-MLD | 0.6 | 0.3 | 0.2 | 0.0 | 0.0 | 100 |
| ΔASF-TLD | 0.6 | 0.3 | 0.2 | 0.0 | 0.6 | 100 |

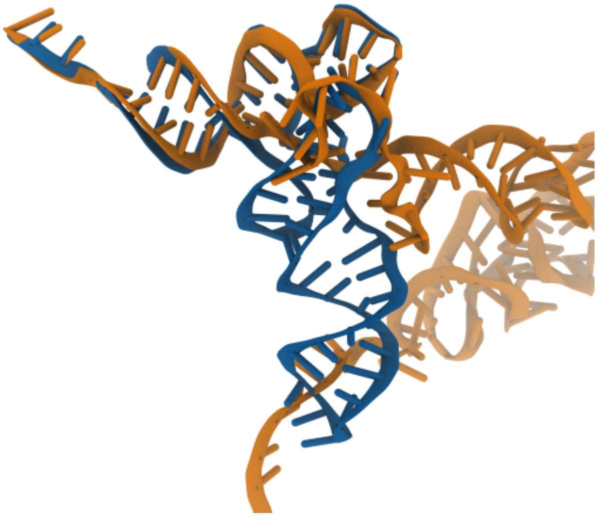

**Figure S1: The tRNA-like domain (TLD) of tmRNA mimics the tRNA acceptor arm** The TLD of tmRNA adopts a structure similar to that of the tRNA acceptor arm. Shown are tRNA in blue (PDB ID: 7AC7[2]) and tmRNA in orange (PDB ID: 7ACJ[2]) bound in the ribosomal P site, superimposed by aligning the large ribosomal subunit (LSU) of the two structures.

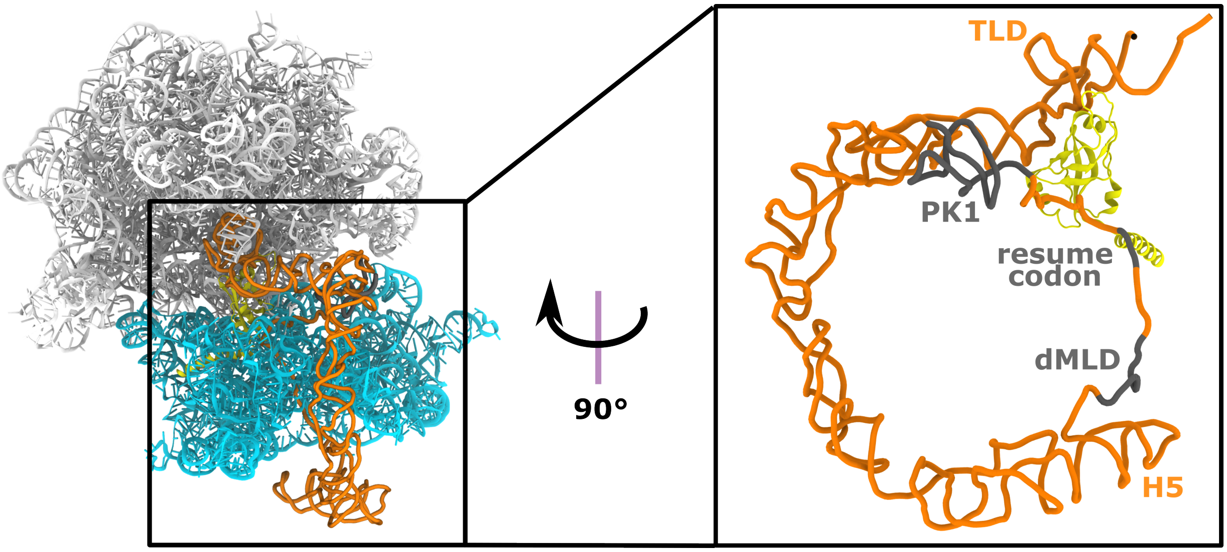


**Figure S2: Structure of tmRNA on the ribosome** (Left) Structure of the ribosomal rRNA. (Right) Rotated and zoomed-in view of the tmRNA, with pseudoknot 1 (PK1) connecting the tRNA-like domain (TLD) with the mRNA-like domain (MLD), which includes the resume codon (G90-A92) and the downstream MLD (dMLD; A97-A103), shown in gray.

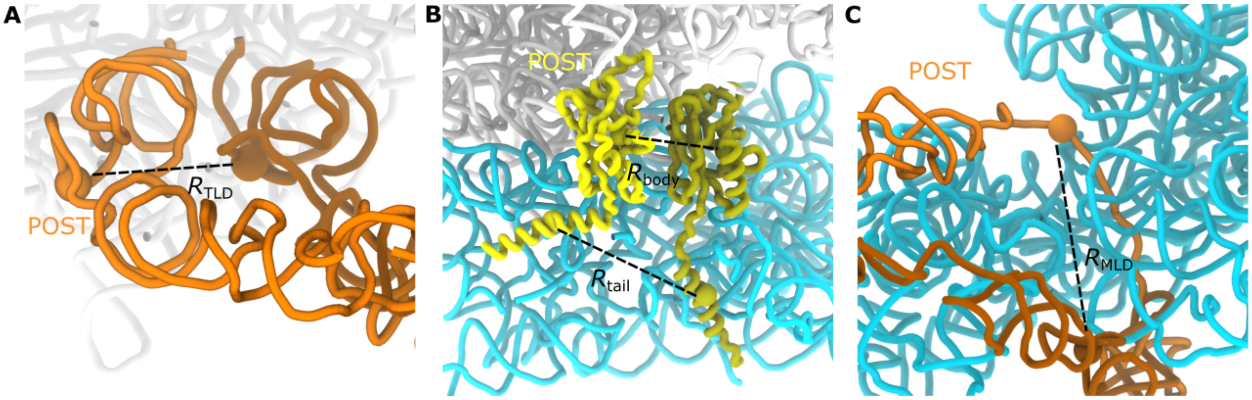


**Figure S3: Coordinate definitions** (A) Definition of $R_{\mathrm{TLD}}$, the distance of the TLD (P atom of U9) from its post-translocation position in the P site, post- and pre-translocation states are shown in light and dark orange, respectively. (B) Definition of $R_{\mathrm{body}}$ and $R_{\mathrm{tail}}^{\mathrm{POST}}$, the distances of the SmpB body ($C_\alpha$ of Leu65) and C-terminal tail ($C_\alpha$ of Glu146) from their post-translocation positions; post- and pre-translocation states are shown in light and dark yellow. (C) Definition of $R_{\mathrm{MLD}}$, the distance of residue G90 (in the resume codon) from its post-translocation position in the mRNA channel; post- and pre-translocation states are shown in light and dark orange.

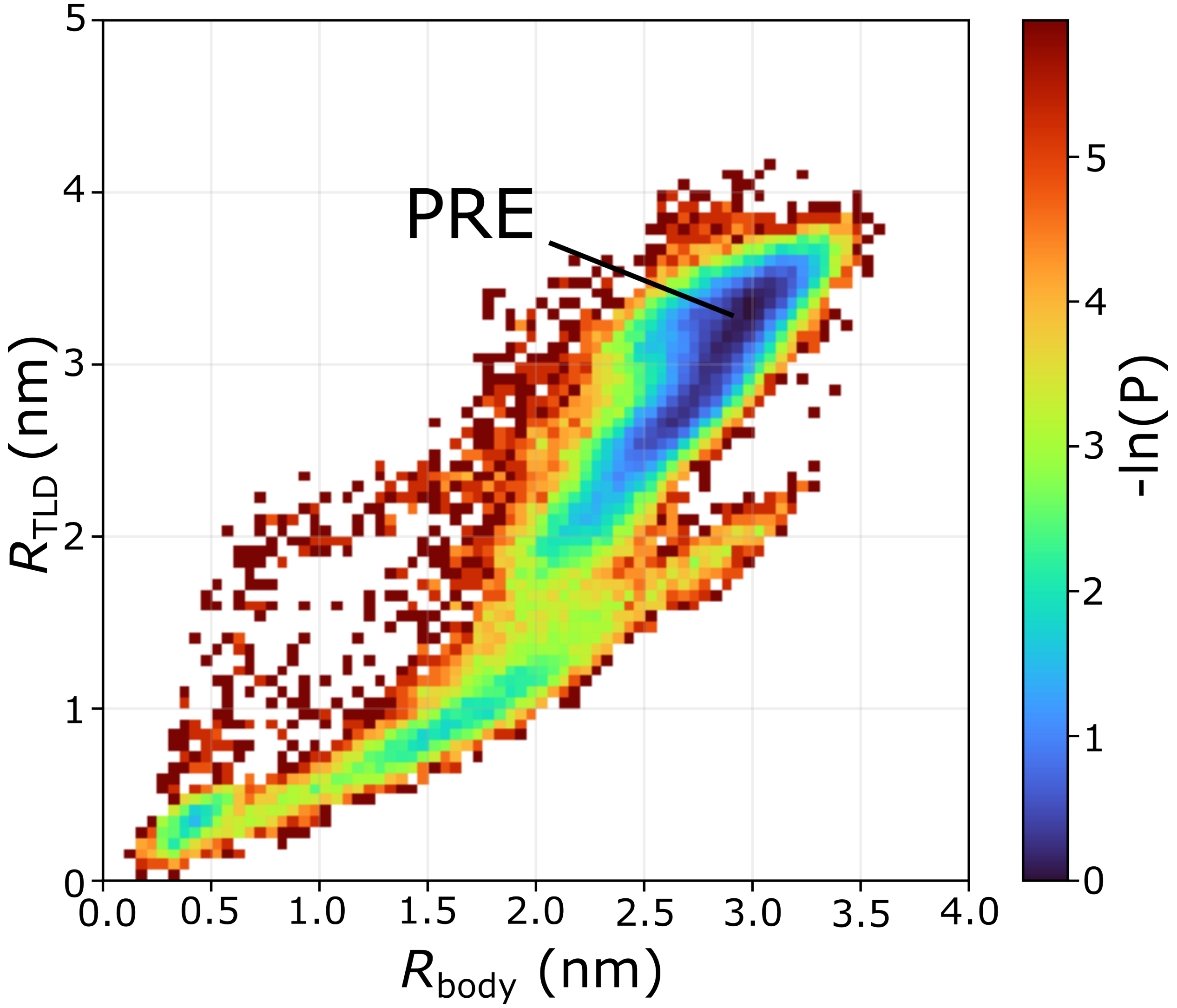


**Figure S4: SmpB body and TLD movements are coupled** Probability distribution as a function of $R_{\mathrm{TLD}}$ and $R_{\mathrm{body}}$. For this distribution, once both coordinates simultaneously reached a value below 0.3 nm, subsequent frames were excluded. This shows that displacement of the SmpB globular domain (the "body") generally follows displacement of the TLD.

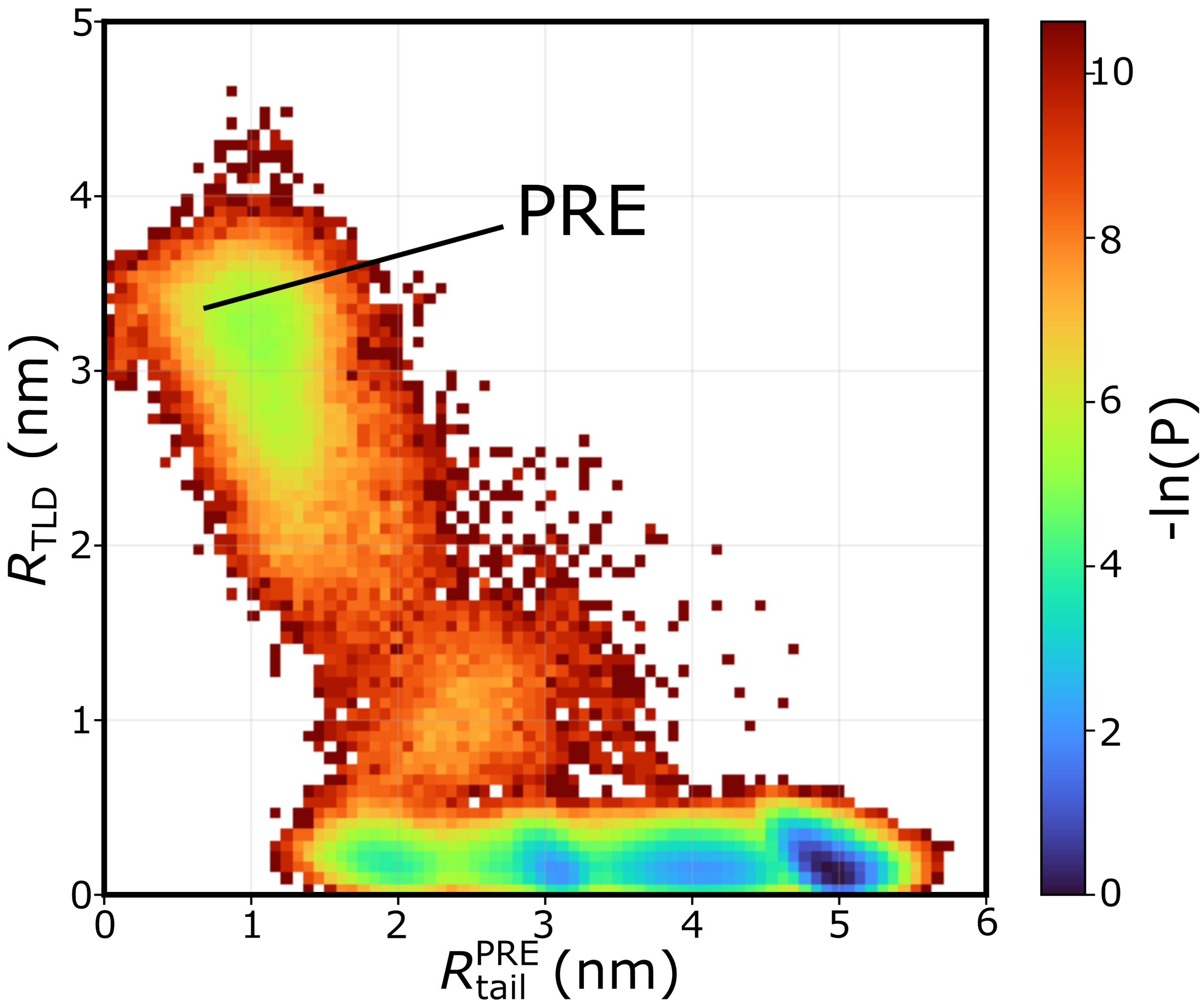


**Figure S5: TLD translocation leads to release of the SmpB tail from the mRNA channel** Two-dimensional distribution of $R_{\mathrm{TLD}}$ and $R_{\mathrm{tail}}^{\mathrm{PRE}}$, illustrating that early in the translocation path the two coordinates are anti-correlated: as $R_{\mathrm{TLD}}$ decreases, reflecting movement of the TLD toward the P site, $R_{\mathrm{tail}}^{\mathrm{PRE}}$ increases, reflecting displacement of the SmpB tail away from its pre-translocation position within the mRNA channel. Once the TLD has fully translocated, $R_{\mathrm{tail}}^{\mathrm{PRE}}$ continues to vary independently over a range of ∼1.5–5.5 nm, indicating there is subsequent rearrangement of the tail.

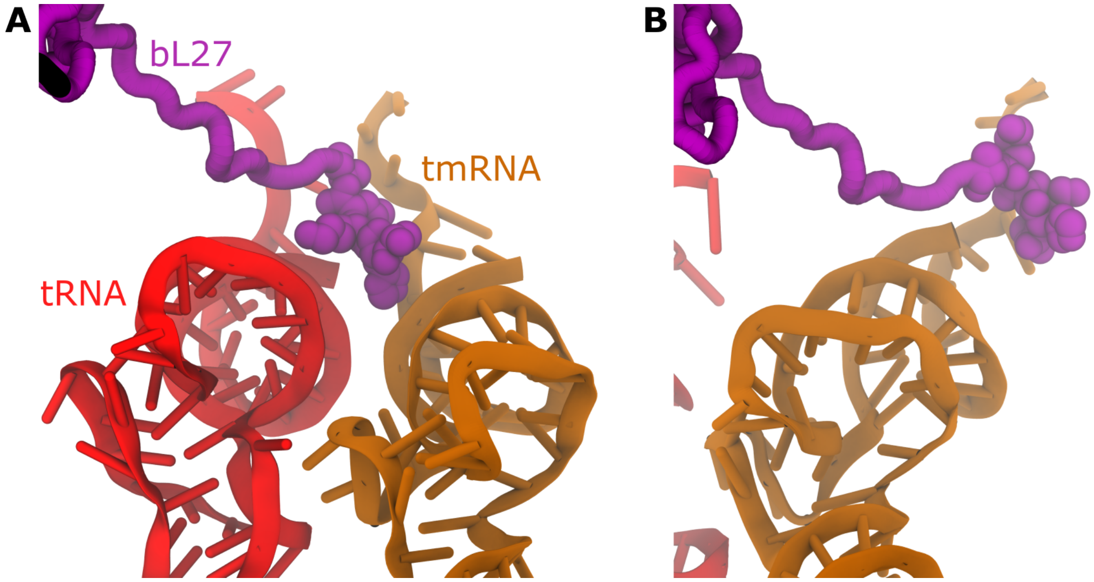


**Figure S6: The N-terminal tail of bL27 impedes TLD translocation** (A) Structure of PRE showing the tRNA (red) positioned in the P site alongside the tmRNA TLD (orange), with bL27 (purple) positioned between the TLD and the P site, thereby obstructing TLD translocation[3]. (B) Structure showing the TLD in the POST state, where it has passed bL27. Together, these structures illustrate how the bL27 N-terminal tail can transiently hinder TLD movement, which would prolong sampling of the accommodated state.

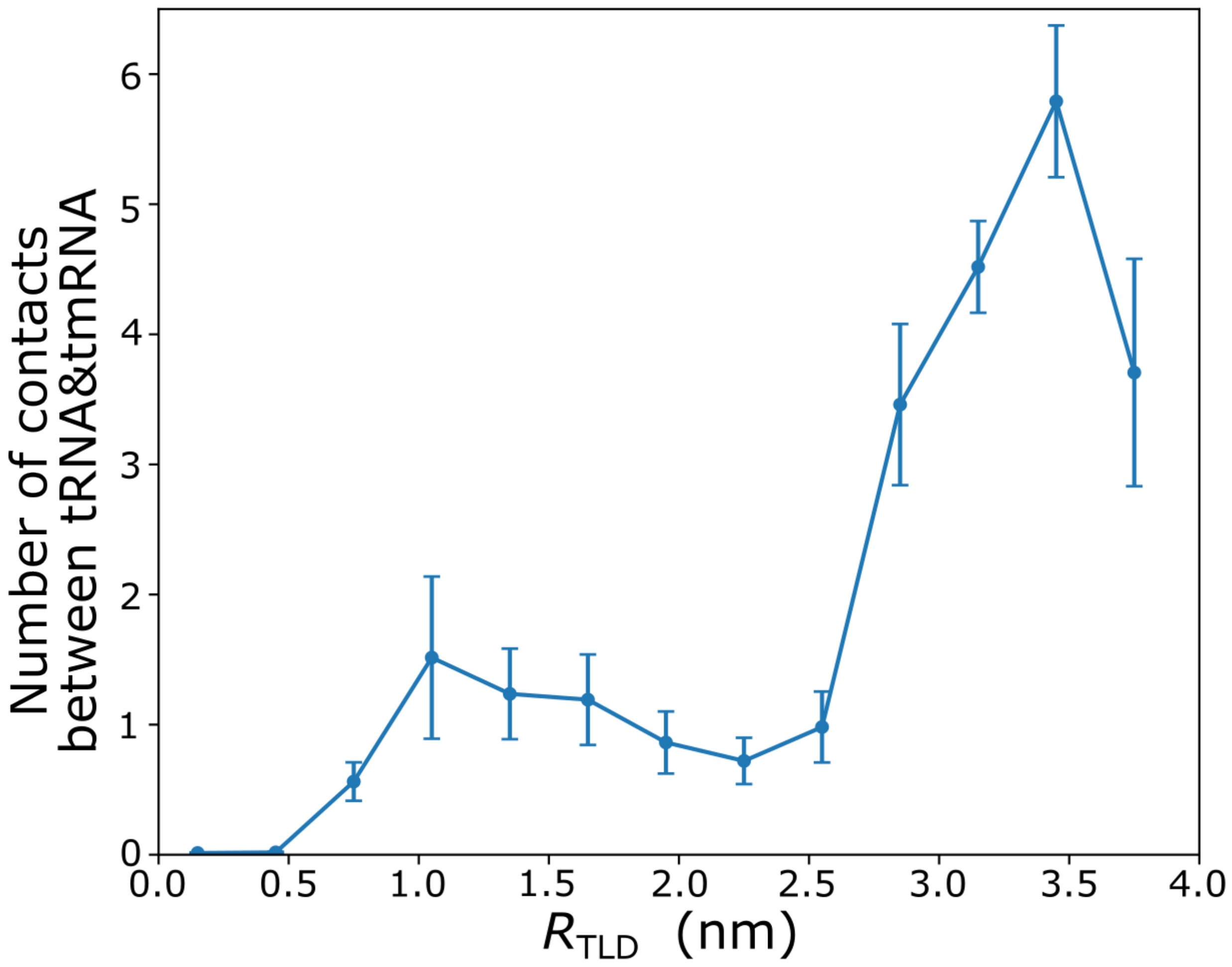


**Figure S7: The P-site tRNA sterically impedes TLD translocation** Mean number of atomic contacts (within 4 Å) between the tmRNA and the P-site tRNA, as a function of $R_{\mathrm{TLD}}$. Between ∼3–4 nm, the TLD maintains more than three contacts with the tRNA, consistent with the P site being physically occluded by the tRNA molecule. Error bars represent the SE across 100 independent runs.

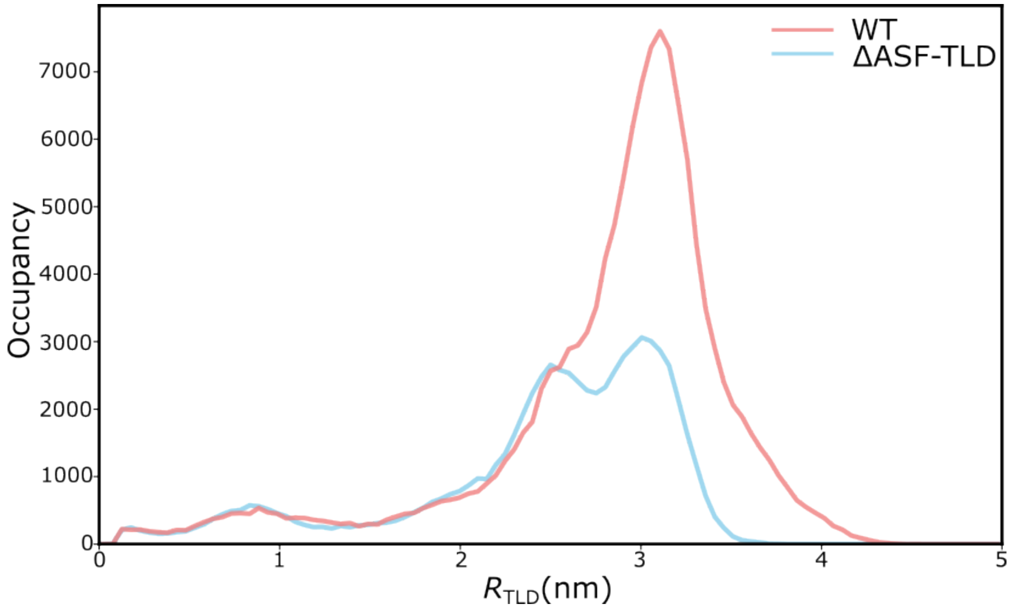


**Figure S8: The A-site finger (ASF) can hinder TLD motion** To test the influence of the ASF on tmRNA dynamics, we generated variants of the model in which repulsive interactions between the ASF and the TLD were removed (ΔASF-TLD; Tab. S1). Occupancy as a function of $R_{\mathrm{TLD}}$ for the Baseline and ΔASF-TLD models. In the Baseline, occupancy is elevated near $R_{\mathrm{TLD}} \approx 3$ nm, corresponding to an accommodated-like state in which the ASF hinders TLD translocation; this occupancy is lower in the ΔASF-TLD model, consistent with relief of the steric barrier.

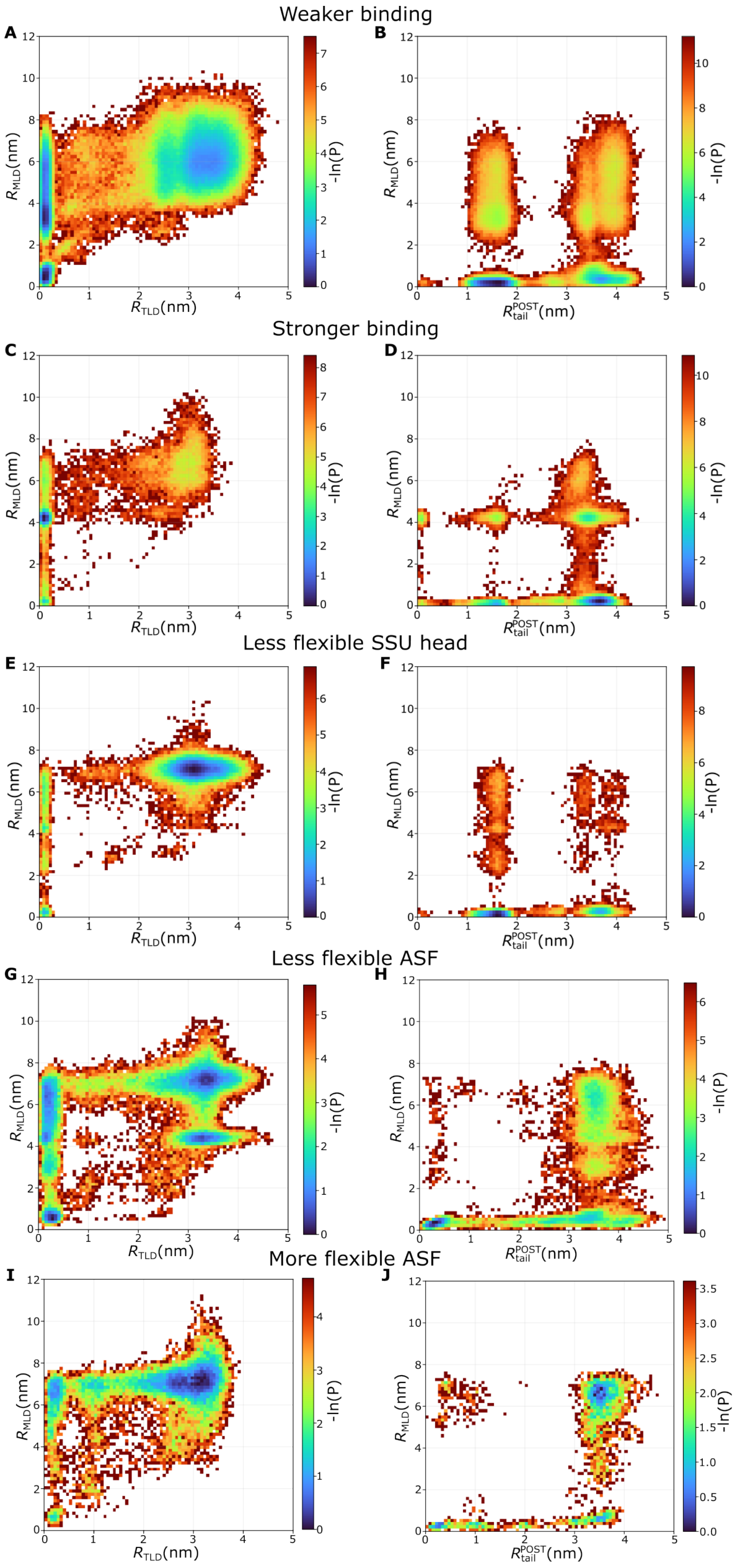


**Figure S9: Robustness of the translocation mechanism across model variants.** To test whether the observed mechanistic features arise from structural rather than energetic factors, we repeated the analysis using several variants of the SMOG model in which the strengths of specific interactions were altered relative to the baseline parameterization (Tab. S1). (A,B) Variant "Weaker binding" in which the contact strength between the tmRNA-SmpB complex and the ribosome in the post-translocation state was scaled by 0.4 (relative to 0.6 in Baseline). Probabilities are shown as functions of $R_{\mathrm{MLD}}$ vs. $R_{\mathrm{TLD}}$ (panel A) and $R_{\mathrm{MLD}}$ vs. $R_{\mathrm{tail}}^{\mathrm{POST}}$ (panel B). (C,D) Variant "Stronger binding", in which this contact strength was instead scaled by 1.0. Probabilities are shown as functions of $R_{\mathrm{MLD}}$ vs. $R_{\mathrm{TLD}}$ (panel C) and $R_{\mathrm{MLD}}$ vs. $R_{\mathrm{tail}}^{\mathrm{POST}}$ (panel D). (E,F) Variant "Less flexible SSU head", in which the flexibility of the SSU head was reduced by scaling the contact strength between the head and the SSU body/LSU to 0.5 (relative to 0.3 in Baseline). Probabilities are shown as functions of $R_{\mathrm{MLD}}$ vs. $R_{\mathrm{TLD}}$ (panel E) and $R_{\mathrm{MLD}}$ vs. $R_{\mathrm{tail}}^{\mathrm{POST}}$ (panel F). (G,H) Variant "Less flexible ASF", in which the flexibility of the A site finger (ASF) was reduced by increasing the strength of the contacts between the SSU and ASF to be scaled by 0.4 (relative to 0.2 in Baseline), Probabilities are shown as functions of $R_{\mathrm{MLD}}$ vs. $R_{\mathrm{TLD}}$ (panel G) and $R_{\mathrm{MLD}}$ vs. $R_{\mathrm{tail}}^{\mathrm{POST}}$ (panel H). (I,J) Variant "More flexible ASF", in which the flexibility of the A site finger (ASF) was increased by reducing the contact strength between the SSU and ASF to be 0.0 (relative to 0.2 in Baseline). Probabilities are shown as functions of $R_{\mathrm{MLD}}$ vs. $R_{\mathrm{TLD}}$ (panel I) and $R_{\mathrm{MLD}}$ vs. $R_{\mathrm{tail}}^{\mathrm{POST}}$ (panel J). In all variants, the overall behavior is preserved, with TLD translocation consistently preceding the other conformational transitions, indicating that this ordering is a robust structural feature.

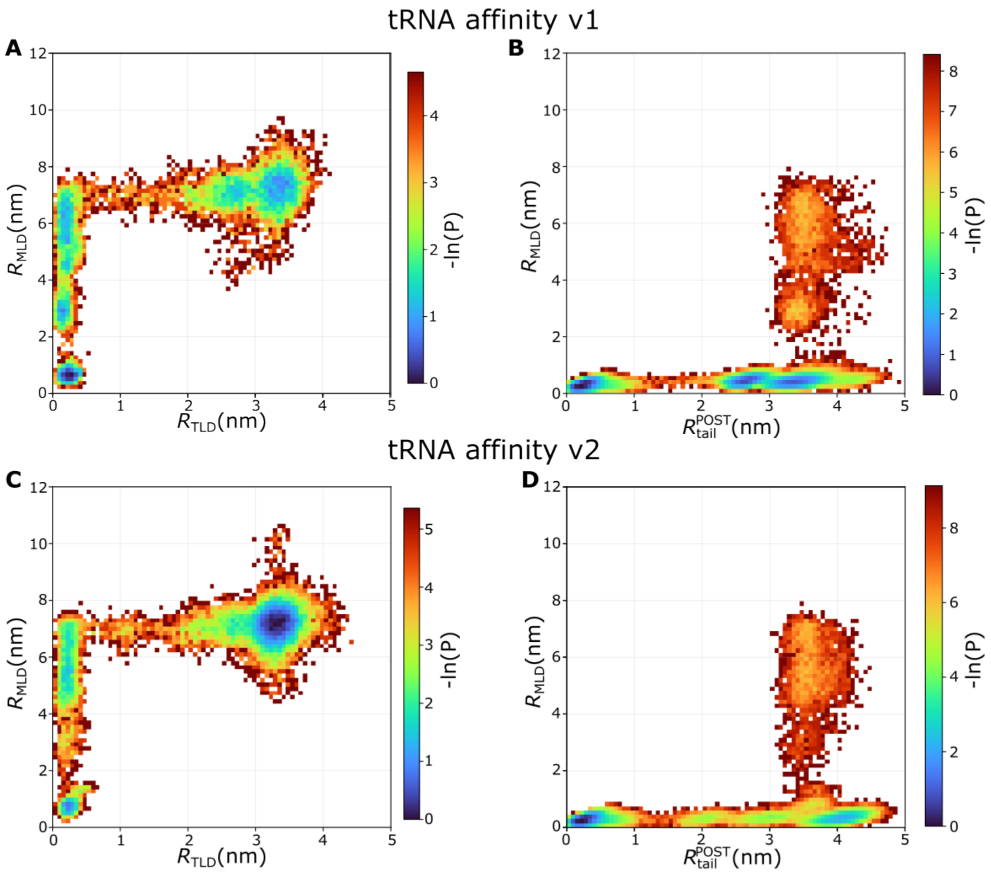


**Figure S10: Robustness of the translocation mechanism to the affinity of tRNA/mRNA for the ribosomal P site.** To further assess the robustness of the mechanism, additional simulations were performed with variants of the model in which the tRNA/mRNA affinity for the P site was altered. This is represented by the contact strength between the tRNA/mRNA and the P site, which was set to 0 in the Baseline model. These stabilizing interactions represents the tighter tRNA/mRNA-P site contacts characteristic of the PRE state. (A,B) tRNA affinity v1 variant with contact strength scaled by 0.1, shown as $R_{\mathrm{MLD}}$ vs. $R_{\mathrm{TLD}}$ (panel A) and $R_{\mathrm{MLD}}$ vs. $R_{\mathrm{tail}}^{\mathrm{POST}}$ (panel B). (C,D) tRNA affinity v2 variant with contact strength scaling increased to 0.2, shown as $R_{\mathrm{MLD}}$ vs. $R_{\mathrm{TLD}}$ (panel C) and $R_{\mathrm{MLD}}$ vs. $R_{\mathrm{tail}}^{\mathrm{POST}}$ (panel D). Despite the shift in the balance between PRE and POST contact weights, the overall behavior is preserved, with TLD translocation consistently preceding the other conformational transitions, confirming that this ordering is a robust feature of the mechanism.

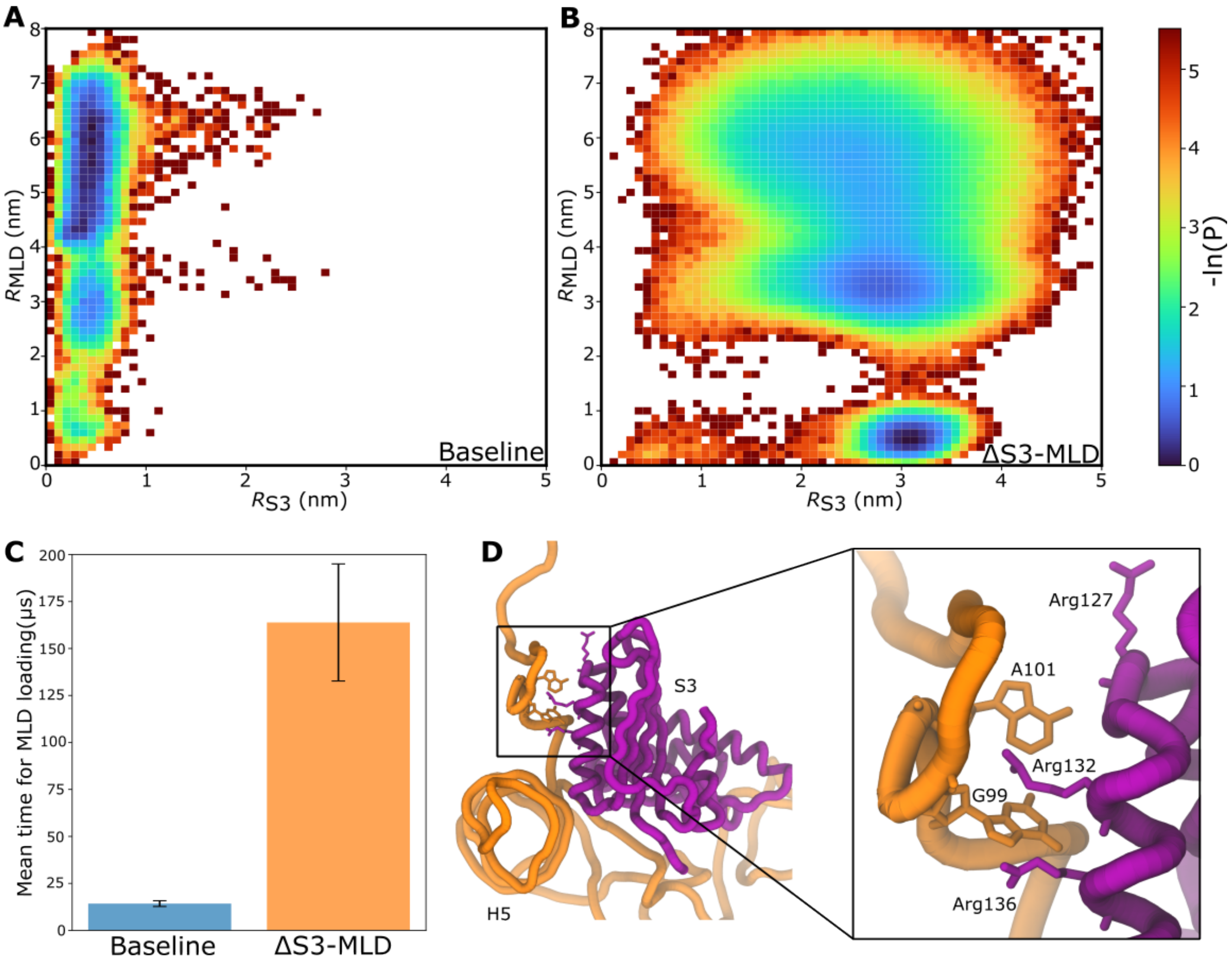


**Figure S11: dMLD–uS3 interactions facilitate MLD loading** (A) Two-dimensional distribution of $R_{\mathrm{MLD}}$ vs. $R_{\mathrm{S3}}$ (the distance of P atom of A100 of the tmRNA from its post-translocation position, reflecting engagement with uS3), shown for frames with $R_{\mathrm{TLD}} < 0.5$ nm. $R_{\mathrm{S3}}$ is already small ($\lesssim$1 nm) across the full range of $R_{\mathrm{MLD}}$ values, indicating that dMLD docking to uS3 occurs prior to and independently of MLD loading, which subsequently proceeds through intermediate states as it fully loads into the mRNA channel. (B) Same as Panel A, shown for the ΔS3-MLD model, in which dMLD-uS3 stabilizing interactions were removed from the force field. Here, $R_{\mathrm{S3}}$ is broadly distributed across all values of $R_{\mathrm{MLD}}$, and MLD loading occurs first, with the overall loading time increased roughly 8-fold relative to Baseline. (C) Mean time for $R_{\mathrm{MLD}}$ to reach $< 0.5$ nm (i.e., fully loaded) for the Baseline and ΔS3-MLD models, computed from 100 independent trajectories per model. The overall loading time increases ∼12-fold for ΔS3-MLD relative to Baseline. Error bars represent the standard error of the mean across trajectories. Because 8 of the 100 ΔS3-MLD trajectories did not reach the loaded state within the simulated time, their time was taken as the last simulated frame for that trajectory; the reported ΔS3-MLD mean is therefore a lower bound on the true loading time. (D) Structural representation of the dMLD-uS3 interface, showing close contacts between dMLD nucleotides G99 and A101 and the positively charged uS3 residues Arg127, Arg132, and Arg136, with Arg132 inserting directly between G99 and A101.

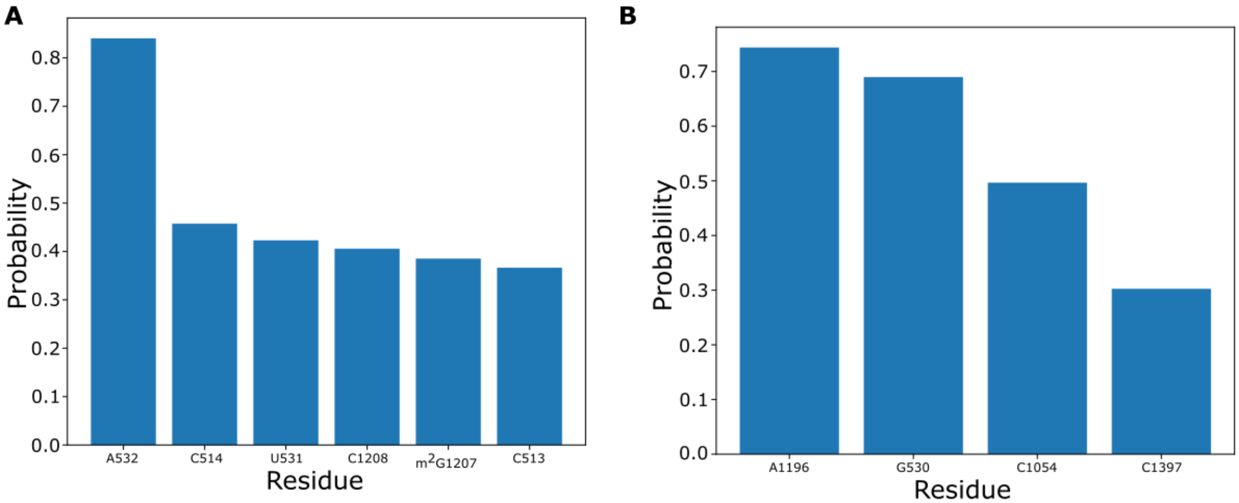


**Figure S12: SSU residues that contact MLD during loading** (A) Probability of SSU residues contacting the MLD when crossing the first barrier ($R_{\text{MLD}}$ between 3.5 and 4 nm ; Figure 3). (B) Probability of SSU residues contacting the MLD during the second barrier ($R_{\text{MLD}}$ between 1 and 2 nm).

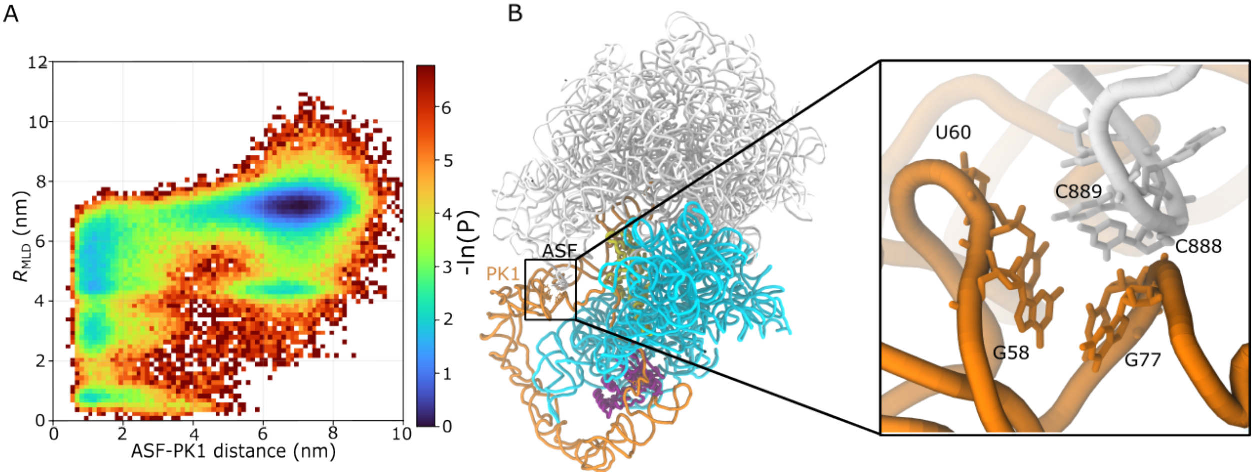


**Figure S13: PK1 forms contacts with the ASF upon TLD translocation.** (A) Two-dimensional probability distribution of MLD loading versus ASF–PK1 distance, showing that along the dominant pathway, the ASF and PK1 come into contact before the MLD is fully loaded. (B) Structural representation showing that, following translocation of the TLD, pseudoknot 1 (PK1) – which connects the TLD to the MLD – comes into contact with the A-site finger (ASF). These interactions with PK1 help stabilize the post-translocation state and, in turn, draw the MLD closer to the small subunit (SSU).

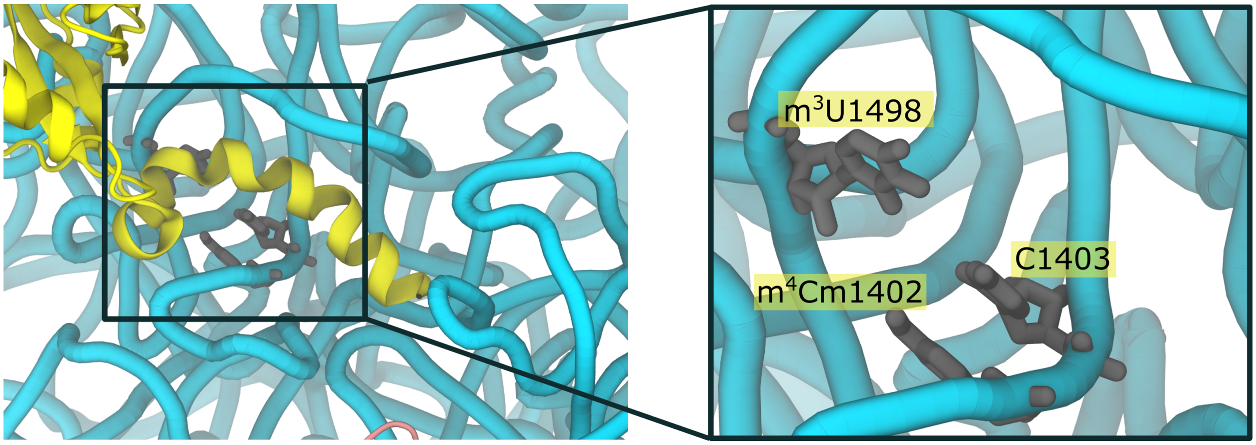


**Figure S14: SSU residues interact with the SmpB tail in the direct intermediate** Structure showing the residues m$^4$Cm1402, C1403, and m$^3$U1498 of the SSU, which interact with the SmpB tail with a probability greater than 0.9 in the direct intermediate, contributing to the steric barrier that lead to this intermediate.

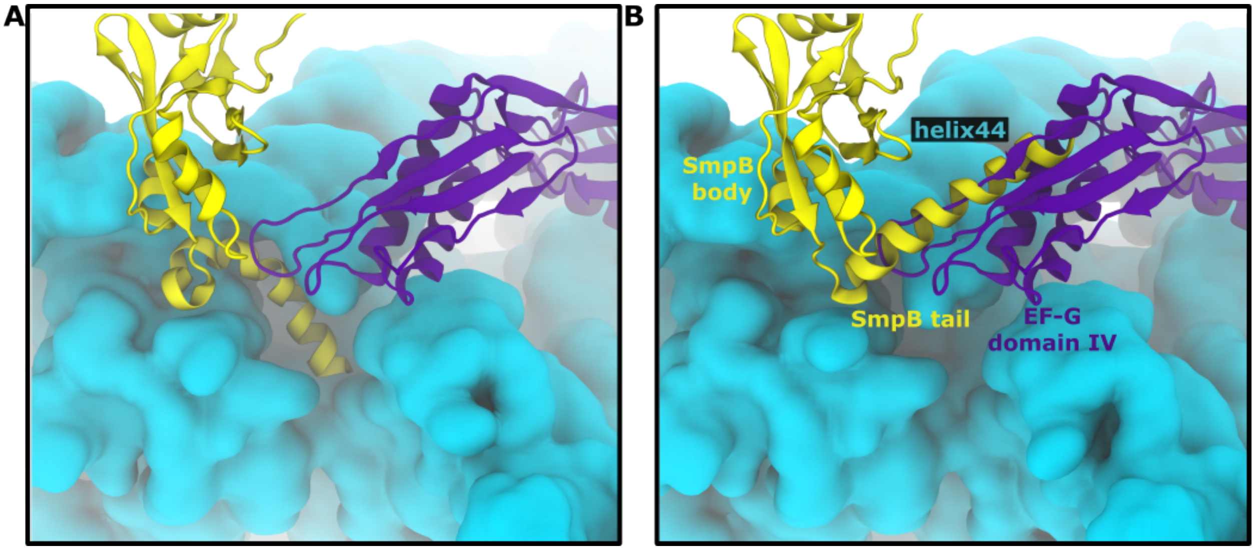


**Figure S15: EF-G overlaps with the position of the SmpB tail in the indirect intermediate** Structures are aligned based on the SSU body in the post-translocation state of tmRNA-SmpB-EF-G complex (PDB 4V6T[4]). (A) In the direct intermediate, the SmpB C-terminal tail occupies a position with minimal overlap with EF-G. (B) In the indirect intermediate, the tail instead occupies a position that almost completely overlaps with domain IV of EF-G (violet) in the vicinity of helix 44. SSU rRNA shown in cyan. This suggests that EF-G may sterically exclude the indirect pathway, preventing the SmpB tail from becoming kinetically trapped.
